# Fungal communities in long-term contaminated soils within a former radium production area

**DOI:** 10.1101/2025.05.05.652156

**Authors:** Anna Rybak, Tatiana Maystrenko, Elena Rasova, Elena Belykh, Ilya Velegzhaninov

## Abstract

The diversity and composition of fungal communities inhabiting soils with technologically increased concentrations of natural radionuclides ^238^U and ^226^Ra within more than 60 years were studied for the first time. Metabarcoding targeting the ITS region was performed using next-generation sequencing technology. Representatives of Basidiomycota and Ascomycota were dominant in both contaminated and reference soils. Additionally, Mortierellomycota and Mucoromycota were subdominant phyla. In general, no significant differences were observed in alpha diversity of soil fungal communities across different contamination levels and seasons. However, when seasonal variations were disregarded, an insignificant trend toward decreased alpha diversity was noted in radionuclide-contaminated soils. Fungi belonging to *Serendipita, Cortinarius, Clavaria, Amphinema, Archaeorhizomyces* were more abundant in the communities of contaminated soils. In contrast, *Thelephora, Tylospora, Russula, Piloderma, Mortierella, Hymenogaster, Hyaloscypha* demonstrated higher abundance in the reference soil. The significant differences in the abundance of certain fungal taxa in the soils of the studied sites mainly involve fungi that are resistant to abiotic and biotic stressors, participate in symbiotic relationships with plant roots and are associated with white rot. These differences likely reflect soil composition and contribute to maintaining the stability of soil fungal communities under conditions of radioactive contamination.

## Introduction

Increased concentrations of natural and artificial radionuclides in the environmental compartments resulting from human nuclear industry activities and catastrophic facility failures (e.g. Chernobyl, Fukushima, etc.) constitute a major anthropogenic threat to both biota and human populations **[1–3]**. Radioactive elements deposited in the soil have a long-term chronic effect on the soil organisms, including fungal and microbial communities **[4,5]**. Fungi play a pivotal role in soil biological processes, influencing soil fertility, decomposition, mineral and organic substance cycling, and plant health and nutrition **[6,7]**. Fungi also contribute to the degradation of toxic organic compounds, including polycyclic aromatic hydrocarbons, pesticides, petroleum hydrocarbons and other hazardous substances **[8–10]**. Certain species of *Aspergillus* and *Trichoderma* can reduce the bioavailability of heavy metals, including radionuclides, by altering the oxidation degree and/or forming insoluble compounds. This process mitigates the risks to living organisms posed by high concentrations of these toxicants in soil **[8–11]**. Also, fungi help reduce risks to biota and the environment associated with the accumulation of metals and radionuclides in soil by chemically modifying these substances or altering their bioavailability **[11]**. Additionally, soil fungi influence the transfer and bioavailability of substances for plants. This suggests that soil fungi, especially those associated with plant roots, could enhance the efficiency of phytoremediation of contaminated soils, including those with elevated concentrations of persistent organic pollutants **[12,13]**. Notably, arbuscular mycorrhizal fungi *Rhizophagus irregularis* isolated from an abandoned uranium mine increased uranium concentrations in the roots and shoots of plants like *Plantago lanceolata* **[14]**.

The capacity of fungi to maintain soil multifunctionality **[7]** and alter the availability of toxicants **[9,15]** renders them excellent candidates for biomonitoring and incorporation in the biological products aimed at restoring technogenically disturbed soils **[16–18]**.

However, most published studies focus on the diversity and composition of fungal communities in soils contaminated with toxic organic compounds or heavy metals **[19–21]**. However, in contrast, published research results on fungal communities in radioactively contaminated territories is limited and often ambiguous. The study **[22]** noted a higher diversity of fungal communities in an area with technologically increased levels of natural radionuclides. The authors believe that changes in the composition and functional structure of microorganisms are directly related to radionuclide and heavy metal contamination, as well as organic matter, and indicate that the community had likely shifted toward a more adapted/tolerant one as evidenced, for example, by the presence of the species Thelephora sp. and Tomentella sp. Also, several 137Cs-tolerant fungal taxa were also identified in fields of 137Cs-contaminated land-use type within a 30-km radius around the FDNPP. At the same time, Qiu and colleagues **[5]** revealed a significant decrease in the richness and diversity of fungal communities in an area contaminated by uranium mining. Also, Portman et al. **[23]** showed lower taxonomic richness and diversity of plant root-associated fungal communities from uranium mine sites than root fungi from paired, off-mine sites in nearby.

In the structure of soil fungal communities inhabiting sites with enhanced concentrations of radionuclides, certain groups resistant to radioactive contamination are observed **[24,25]**. Conversely, the abundances of several groups were significantly reduced in soil samples due to uranium mining activities **[5]** or exposure to gamma irradiation **[26]**. In the study of the microbial communities in a former pilot-scale uranium mine in Eastern Finland **[25]** the authors found that the fungal genera *Archaeorhizomyces, Oidiodendron*, unclassified Trichocomaceae, *Hygrocybe* and *Geminibasidium* correlated negatively and significantly with ^226^Ra. The relative abundance of unclassified fungi also correlated negatively and significantly with ^226^Ra. In contrast, *Pseudeurotum*, Holwaya, *Tremellomycetes* (unclassified) and *Cryptococcus* had significant positive correlation with ^226^Ra.

Representatives of this kingdom exhibit resistance to increased levels of ^137^Cs in soils from areas contaminated by nuclear disasters **[13,27]** as well as in controlled laboratory experiments **[28]**. One of approaches to enhancing fungal resistance to ionizing radiation is to increase their melanin content **[29–31]** oth increases **[25]** and decreases in community diversity, as well as changes in community structure **[5]**.

The aim of our investigation was to examine the diversity and structure of fungal communities in soils contaminated with naturally occurring radionuclides for over 60 years using metabarcoding targeting the ITS region. Analyzing fungal communities that have developed and function in technogenically disturbed radioactive territories provides a new knowledge on the resistance of fungal groups to extreme exposure and serves as a foundation for developing and refining bioremediation technologies in radioactively contaminated areas.

## Materials and Methods

### 2.1. Study area and soil characteristics

The investigation was conducted on the territory of a former radium production facility with increased levels of ^238^U и ^226^Ra in soil around Vodny settlement (Republic of Komi, Russia) and a nearby reference site. Radioactive contamination in the area resulted from the operations of radiochemical plants, which extracted ^226^Ra from highly mineralized groundwater and recycled uranium waste during the 1930s to 1960s. Furthermore, the site contaminated with radioactive materials was also used for producing charcoal for these radiochemical plants. So, this territory contaminated with heavy natural radionuclides serves as a natural testing ground for studying the effects of chronic ionizing radiation on biota.

Soil samples were collected in August and October 2023 (10 samples per site in total), flash-frozen in liquid nitrogen and stored at −80 °C until analysis. The vegetation at both sites is characterized by mixed forest on Gleyic Retisols **[32]**. The level of γ-radiation background (μSv/h) before sampling was measured using a DKG02U “Arbitr” dosimeter (SPC “Doza”, Russia) at a height of 1 m from the soil surface. The pH values of the salt (1M KCl) extract were determined by potentiometry using pH-meter pH-410 (Aquilon, Russia), total carbon - by gas chromatography on the element analyzer EA 1110 (CHNS-O). The granulometric composition of the soil was determined using the laser diffraction method with the particle size analyzer Bettersizer 2600 (China) coupled with an ultrasonic particle disperser “UZDN-1” (energy of 450 J/cm3) **[33]**. The chemical analyses of the soils were conducted in the “Ecoanalit” Laboratory and the Laboratory of Radionuclide Migration and Radiochemistry of the FRC Komi SC UB RAS. A detailed description of the methods used to measure radionuclide activity concentrations and other procedures of chemical analysis are provided in **[34,35]**.

The level of the γ-radiation background ranged from 1.5 to 8.7 μSv/h at the contaminated site, and from 0.1 to 0.5 μSv/h at the reference site. In soils of the contaminated site, the concentrations of ^238^U and ^226^Ra were 241–3629 Bq/kg and 9–238 kBq/kg, respectively. In contrast, soils of the reference site had concentrations of 11–254 Bq/kg for ^238^U and 0.1–2.9 kBq/kg for ^226^Ra. Notably, during the production of ^226^Ra, coal derived from wood burning was used in technological processes, resulting in high charcoal concentrations in all soil samples from the contaminated site. The total carbon content in soils of the contaminated site ranged from 3.2 to 12.6 %, while in the reference soils from 1.1 to 9.2 %. Soil pH values varied from 4.8 to 6.8 at the contaminated site, and from 3.4 to 5.9 at the reference site. The physical composition of the soils consisted of 75% sand and 25% clay at the contaminated site, compared to 85% and 16%, respectively, at the reference site.

### 2.2. Metagenomic analysis

Total DNA was extracted from 500 mg of soil using HiPure Soil DNA Kit with silica gel columns (Magen, China) and additionally purified using ColGen kit (Syntol, Russia) following the manufacturer’s protocols. All procedures for DNA isolation and purification were performed at the Center for Collective Use «Molecular Biology» of the IB FRC Komi SC UB RAS. DNA concentration was measured using <u>Quantus™ Fluorometer</u> (Promega, USA) with the QuantiFluor® ONE dsDNA System. The ITS2 region was amplified using the primer pairs ITS3_KYO2/ITS4 (forward: 5′-GATGAAGAACGYAGYRAA-3′, reverse: 5′-TCCTCCGCTTATTGATATGC-3′), combined with Illumina adapter sequences. Amplicon libraries were prepared according to the following steps: 1) PCR amplification: 95°C for 3 min, followed by 25 cycles of 95°C for 30 s, 55°C for 30 s, 72°C for 30 s and a final elongation of 72°C for 5 min; 2) PCR purification using AMPure XP Beads particles; 3) index amplification; 4) PCR purification using AMPure XP Beads particles; 5) evaluation of the quality of the prepared products and measurement of concentrations; 6) normalization and pooling of prepared libraries; 7) libraries were cleaned and mixed equimolar using SequalPrep™ Normalization Plate Kit (Thermo Fisher Scientific, USA); 8) quality control of the received library pools was carried out using the system Fragment Analyzer. Amplicon libraries were sequenced using 2×250 bp paired-end reagents on a MiSeq platform (Illumina, USA).

### 2.3. Bioinformatic and statistical analysis

All statistical analyses were performed using R software version 4.4.1 **[36]**. Sequence processing was conducted using the *dada2* package following an ITS-specific version of the DADA2 tutorial workflow **[37]**. Amplicon Sequence Variant (ASV) sequences were compared against the UNITE ITS database for fungi (release date 04/04/2024) **[38]**. Raw sequences have been deposited in the NCBI Sequence Read Archive under BioProject accession number PRJNA1154587. ASV diversity and richness were analyzed using the *phyloseq* and the *vegan* packages **[39,40]**, after filtering and rarefying the data. Differential abundance analysis of ASV was performed using *DESeq2* **[41]**. The Wilcoxon test was used to assess differences in fungal community diversity parameters between the studied sites.

## Results

### 3.1. Alpha diversity

A range of diversity metrics were employed to assess the alpha diversity of soil fungal communities. Higher diversity values were observed in the fungal communities of the contaminated site compared to the reference site during the summer period (Table 1).

**Table 1.** Alpha diversity metrics of soil fungal communities at different sites (M ± SD).

| Season | Site | Observed | Shannon | Chao1 | ACE | Simpson | Fisher |
| --- | --- | --- | --- | --- | --- | --- | --- |
| Summer | Contaminated | 166.6 ± 15.1 | 3.6 ± 0.2 | 166.8 ± 15.1 | 167.1 ± 15.1 | 0.94 ± 0.02 | 26.8 ± 2.6 |
|  | Reference | 161.2 ± 36.0 | 3.4 ± 0.5 | 161.5 ± 36.4 | 161.5 ± 36.3 | 0.92 ± 0.04 | 25.9 ± 6.2 |
| Autumn | Contaminated | 122.2 ± 26.4 | 2.9 ± 0.5 | 123.4 ± 25.6 | 122.5 ± 26.3 | 0.87 ± 0.08 | 19.6 ± 4.7 |
|  | Reference | 148.6 ± 34.8 | 3.2 ± 0.6 | 148.9 ± 34.7 | 149.1 ± 34.6 | 0.89 ± 0.06 | 23.5 ± 6.3 |
| Combined data | Contaminated | 144.4 ± 31.4 | 3.3 ± 0.5 | 145.1 ± 30.7 | 144.8 ± 31.5 | 0.90 ± 0.05 | 21.0 ± 5.3 |
|  | Reference | 154.9 ± 8.9 | 3.3 ± 0.1 | 155.2 ± 8.9 | 155.3 ± 8.8 | 0.90 ± 0.02 | 22.8 ± 1.5 |

Conversely, in autumn, the fungal communities at the contaminated site were less diverse than those at the reference site. The increased sensitivity of the fungal community at the contaminated site to seasonal changes may be attributed to the high charcoal concentrations in the soil and the diverse biochemical processes associated with this material, involving various microbial groups within the studied community. Overall, the diversity of fungal communities at both sites was lower in autumn than in summer. However, all differences were found to be statistically insignificant.

When analyzing the aforementioned parameters without considering seasonal variations, no statistically significant differences were found between the alpha diversity indices of the radioactively contaminated and reference soils. In autumn, the alpha diversity indices, such as Chao1 and ACE, were higher at the reference site compared to the contaminated site, although these differences were statistically insignificant (Table 1).

### 3.2. Soil fungal community composition

ASV related to 15 phyla, 114 orders, 47 classes, 206 families, 278 genera and 115 species were identified in soil fungal communities of both contaminated and reference sites during summer and autumn. Basidiomycota (38.3-82.9%) and Ascomycota (13.0-64.1%) were dominant phyla in all analyzed samples, with a relative abundance >1%. Subdominant phyla included Mortierellomycota (up to 6.0%) and Mucoromycota (up to 6.6%) (Figure 1).

**Figure 1.**
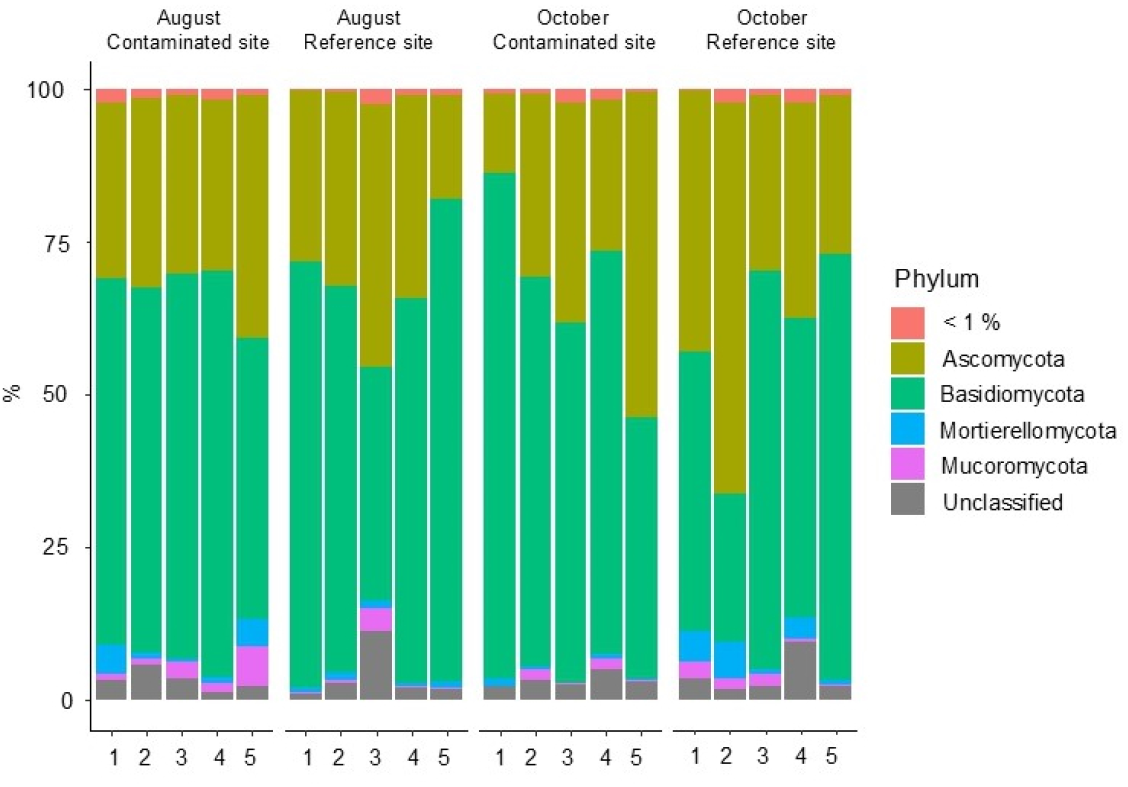
The phylum composition of fungal communities with a relative abundance >1% in the soils of the investigated sites varied across different seasons.

The combined share of phyla with a relative abundance <1% including Rozellomycota, Glomeromycota, Neocallimastigomycota, Chytridiomycota, Olpidiomycota, Zoopagomycota, Calcarisporiellomycota, Blastocladiomycota, Sanchytriomycota, Kickxellomycota, and GS01_phy_Incertae_sedis varied from 0.1 to 2.4 %. The abundance of the unclassified fungi (Unclassified in Figure 1) ranged from 1.1 to 11.3 % across all samples.

At the family level Cortinariaceae and Russulaceae dominated across all analyzed samples (Figure 2).

**Figure 2.**
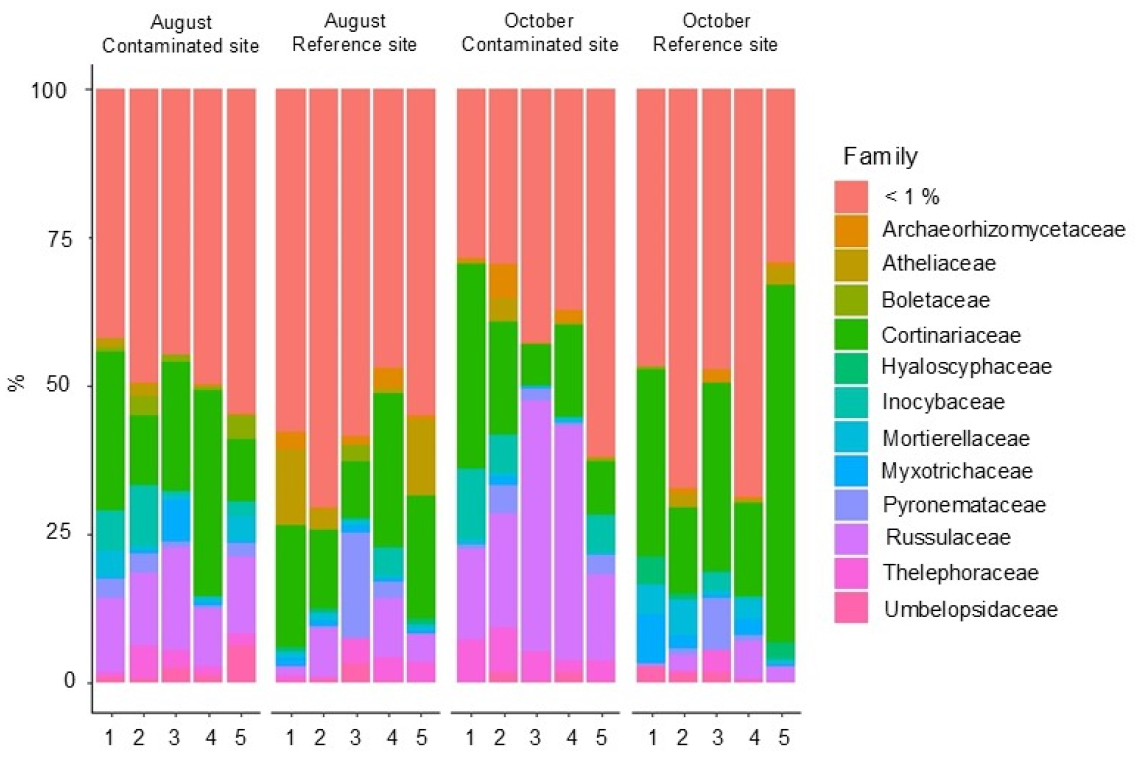
The family composition of fungal communities with a relative abundance >1% in the soils of the investigated sites varied across different seasons.

The relative abundance of ASVs related to Cortinariaceae ranged from 7.0 to 60.1 %. Representatives of Russulaceae were identified in soils from both sites with abundances ranging from 0.17 to 42.3 %. The remaining families with a relative abundance >1% are presented in Figure 2.

At genus level it was shown that the majority of ASVs had a relative abundance of less than 1% (30.8-81.0%) The most abundant ASVs classified at the genus level were *Cortinarius* (7.0-60.1%), *Russula* (0.2-41.9%), *Mortierella* (0.1-4.8%), *Umbelopsis* (0.1-6.1%). Other notable ASVs thogh not consistently identified across all soil samples included *Wilcoxina* (0.1-17.7%), *Inocybe* (0.1-11.6%), *Tomentella* (0.02-6.8%), *Leccinum* (0.03-4.0%), *Oidiodendron* (0.1-8.4%) and *Archaeorhizomyces* (0.1-5.8%) (Figure 3).

**Figure 3.**
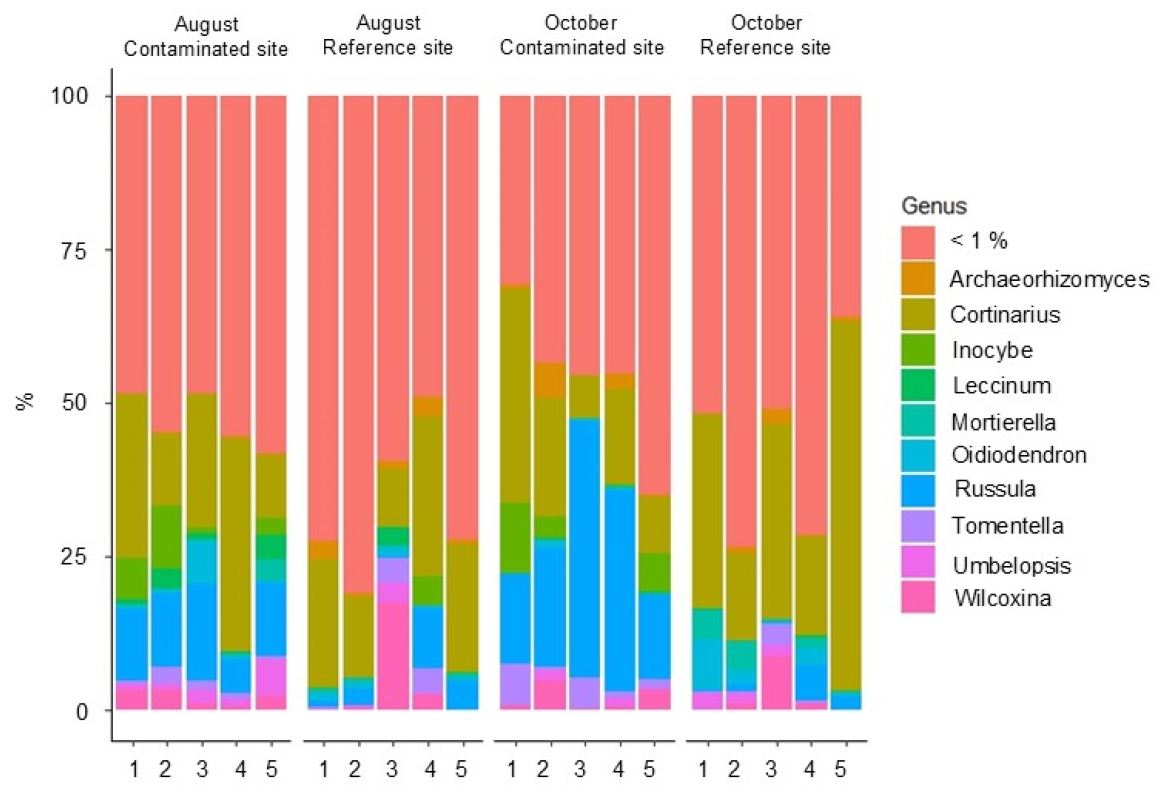
The genus composition of fungal communities with a relative abundance >1% in the soils of the investigated sites varied across different seasons.

### 3.3. ASV differential abundance analysis using DESeq2

In our investigation the differences in ASV abundance between the soil fungal communities of the contaminated and reference sites were analyzed. The results of this assessment at the phylum, family and genus levels are presented in Figure 4.

**Figure 4.**
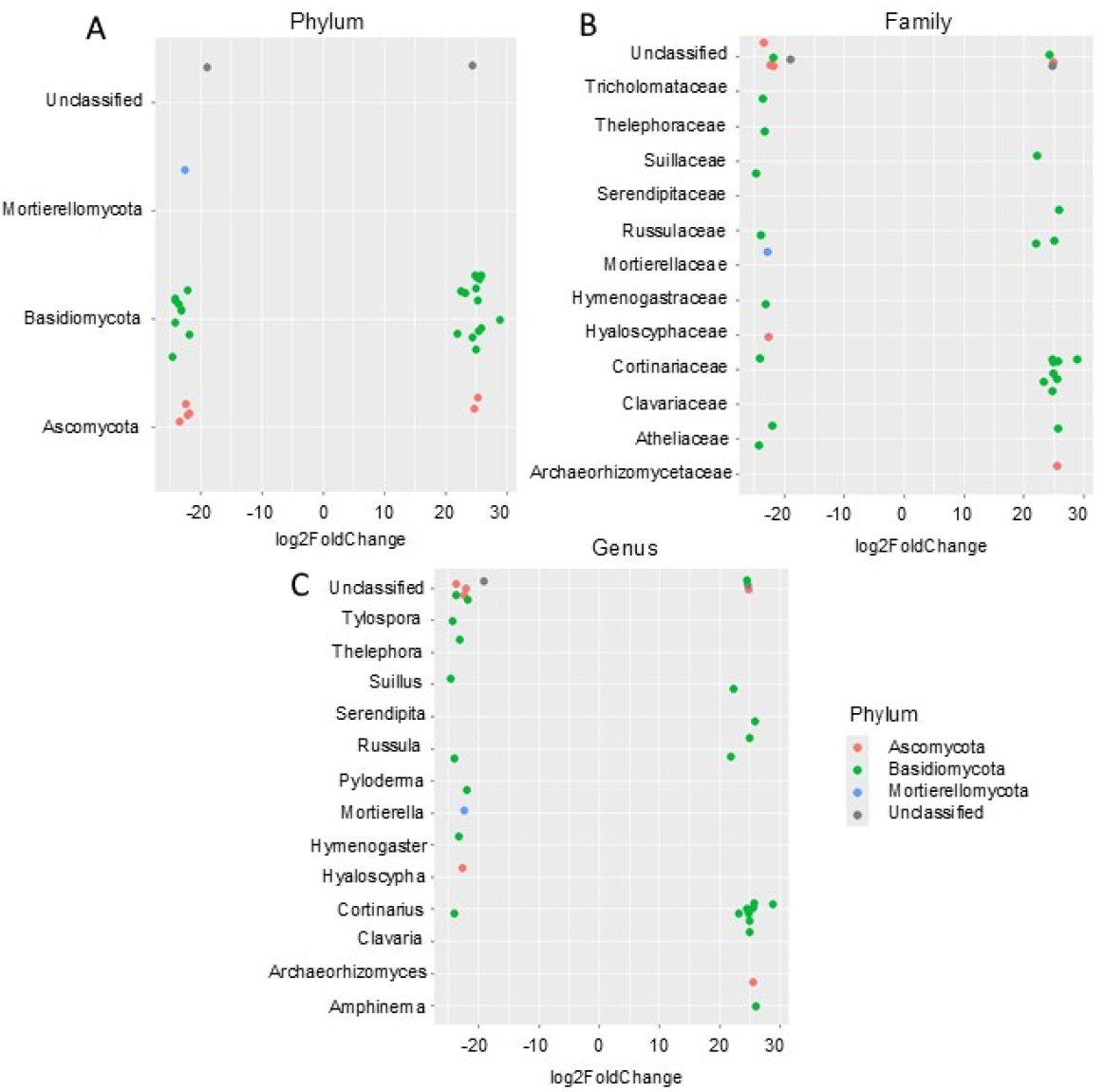
Significant differences in ASV abundances between the contaminated and reference sites identified at the phylum (A), family (B) and genus (C) levels using DESeq2. The figures represent log2 fold changes in ASV abundances between the sites: below 0 were more enriched in the group of reference samples, while those with values above 0 were more enriched in the group of contaminated samples.

The significance of the differences in ASV abundances between the contaminated and reference sites is detailed in Table S1 in Supplementary. Significant differences were observed for 32 ASVs identified in the fungal communities of the contaminated and reference soils. At the phylum level Basidiomycota was the most abundant with the majority (22 from 32) of significantly differing ASVs belonging to this phylum between the reference and contaminated sites. Additionally, several ASV from Ascomycota and one from Mortierellomycota were also identified.

At the lower taxonomic level among Basidiomycota, Cortinariaceae was the most diverse, with eight representatives of the *Cortinarius* genus. Six of these ASVs exhibited significantly higher abundance at the contaminated site: ASV15 (logFC = 29, *p* = 1.17E-27), ASV17 (logFC = 23, *p* = 3.51E-15), ASV21 (logFC = 25, *p* = 2.16E-17), ASV83 (logFC = 26, *p* = 6.51E-18), ASV88 (logFC = 26, *p* = 4.26E-18), ASV117 (logFC = 25, *p* = 4.46E-17), ASV124 (logFC = 25, *p* = 5.81E-17). Two *Russula* sequences (ASV13 and ASV207) showed higher abundances at the contaminated site (logFC > 21, *p* = 1.91E-17 and *p* = 1.38E-13, respectively) while one (ASV43) was more abundant at the reference site (logFC = -24, *p* = 8.06E-19). ASVs belonging to *Suillus* were found in soils from both sites: ASV63 was significantly more abundant in the reference soil (logFC = 25, *p* = 5.32E-17), whereas ASV109 was more abundant in the contaminated soil (logFC = 5, *p* = 4.21E-14). The abundance of ASVs identified as *Serendipita* (*p =* 1.80E-18), *Clavaria* (*p =* 3.87E-17), *Amphinema* (*p =* 2.45E-18) from Basidiomycota and *Archaeorhizomyces* (*p =* 7.40E-20) from Ascomycota was higher at the contaminated site (logFC > 25 for all these ASVs) (Figure 4). Conversely, representatives of Basidiomycota (*Thelephora, p =* 3.93E-15; *Tylospora, p =* 2.34E-16; *Russula, p =* 8.06E-19; *Piloderma, p =* 9.49E-1*4*; *Hymenogaster, p =* 3.87E-15), Ascomycota (*Hyaloscypha, p =* 2.60E-14) and Mortierellomycota (*Mortierella, p =* 2.24E-14) showed pronounced decreases (logFC > -22 for all these ASVs) in abundance in the contaminated soil compared to the reference soil.

## Discussion

Our research performed using ITS metabarcoding has provided new insights into how fungal communities respond to radioactive contamination of the soil. The data we have obtained on the diversity, composition, and structure of soil fungal communities containing elevated concentrations of natural radionuclides complement the limited information provided in the published works. In the future, certain taxa identified in the fungal communities of radioactively contaminated soils can be used in the development of technologies and approaches for bioremediation of technogenically disturbed territories.

Alpha diversity, which reflects the variety within a community, primarily depends on the number of identified taxa. In general, without considering for seasonal variations, there was a slight trend towards decreased alpha diversity in radionuclide contaminated soil; however, no significant differences were observed (Table 1, Combined data). Studies investigating soil fungal communities in areas with high concentrations of naturally occurring radionuclides have yielded ambiguous results. On the one hand some studies have reported a decrease **[5]** in richness and diversity indicators of soil fungal communities or even the near-complete destruction **[42]** of fungal communities. On the other hand, other investigations have observed high diversity of soil fungi **[22,25]**. These contradictions in the data may be explained by differences in the composition of co-contaminants or variations in plant community structure. So, results of the study **[43]** of the microbial community in an arid habitat with radiation stress (^137^Cs) shown, that the highest diversity of the fungal community was in spring. And season was a determinant factor in the root endophyte community composition.

It should also be said that probably, the introduction of coal into the soil during radium production significantly reduces the thermal conductivity of the soil **[44,45]**, which clearly affects the rate of freezing and the crystallization process of water. Additionally, coal influences nutrient exchange **[46]** and gas exchange **[47]** in the soil, which in turn may have an effect on the survival and reproduction of certain fungal groups during seasonal temperature and humidity changes and in freeze-thaw cycles.

Basidiomycota and Ascomycota dominate the fungal community structure in the soils of the studied area, while Mortierellomycota and Mucoromycota occupy subdominant positions. This community structure is characteristic of soil fungal communities of high-latitude boreal forests **[48,49]**. Similar patterns have also been observed in mine-contaminated soils of highlands **[50]**. The structure of soil fungal communities in an area referenced as oil-contaminated **[51,52]**, located approximately 500 km northeast of the radioactive-contaminated territory under study, showed a dominance of Ascomycota over Basidiomycota. Subdominant positions were occupied by Zygomycota, Glomeromycota, and Rozellomycota **[51]** or by Zygomycota and Chytridiomycota **[52]**. Overall, the distribution of dominant fungal phyla at the reference and oil-contaminated sites was similar. However, the location at the border of the same soil and climate zone may be a more significant factor influencing the composition of soil fungal communities. This should be considered when comparing results from different research groups.

A high number of fungi are known to have symbiotic interaction with both high plants and the liverworts **[53]**. Mycorrhizal associations promote ecosystem restoration by facilitating plant communities’ development **[54]** *de novo* and after natural disasters. Representatives of the Basidiomycota and Ascomycota phyla participate in predominately ectomycorrhizal associations **[55]** and wood decomposition **[56,57]**. Basidiomycota are known for their specialized ability to degrade plant polysaccharides and decompose lignin. In contrast, fungi from Ascomycota predominantly metabolize crop residues and litter, focusing on polysaccharides rather than lignin as their primary substrate **[56]**.

The distribution of lower taxa within dominant phyla is influenced by specific environmental conditions. The most significant changes in the structure of fungal communities were observed at lower taxonomic levels such as family and genus. Using DeSeq2 analysis our study identified groups specific to the soils of both the contaminated and reference sites. At the genus level, specific ASVs belonging to *Serendipita, Clavaria, Amphinema, Archaeorhizomyces* were more abundant in fungal communities of the contaminated soil (Figure 4C). Conversely, ASVs from genera such as *Thelephora, Tylospora, Russula, Piloderma, Mortierella, Hymenogaster, Hyaloscypha* were the most abundant in soils of the reference site. Among the genera mentioned were fungi capable of forming ectomycorrhizal associations (*Amphinema:* **[58]**, *Archaeorhizomyces:* **[59]**, *Russula:* **[60]**, *Tylospora:* **[61]**, *Piloderma*: **[62,63]**, *Suillus:* **[64,65]**, *Hymenogaster* and *Thelephora*: **[66]**).

Ectomycorrhizal associations are commonly found in temperate zone **[67]**, and enhance tolerance to harsher conditions **[68,69]**. Therefore, the detection of these fungi within the studied fungal communities is anticipated. It should also be noted that members of the genus *Serendipita*, which are abundant in the contaminated site soils, are recognized as fungal endophytic symbionts that enhance plant resistance to abiotic and biotic stress **[70,71]**. As one of the best endophyte fungi *Serendipita indica* (*Piriformospora indica*) was shown **[72]**. Because in addition to reducing heavy metal stress for crops this taxon could significantly reduce the threat of other abiotic stresses. In particle, *Serendipita indica* enhance the growth and productivity of plants in contaminated environments by improving soil quality, reducing cadmium absorption, improving the activity of antioxidant enzymes and secondary metabolites, raising water and mineral absorption, and altering morphophysiological structures. Also, *Serendipita* and *Cortinarius* abundancies were found to increase together with the soil carbon content increase as a result of forest fires **[73]**.

Interestingly, seven ASVs belonging to the genus *Cortinarius* were identified among the dominant taxa in the contaminated soil samples. Additionally, a high abundance of *Hebeloma* was recorded in three samples from this site. These fungi, classified as white rot species, are capable of degrading wood, particularly lignin, without causing acidification of the environment, unlike brown rot species **[57]**. This phenomenon is likely attributable to the large amount of charcoal present in the soil at the contaminated site.

Overall, the observed microbial diversity reflects environmental conditions and the dominant taxa represent the most adapted forms **[74]**. However, to date, data on the composition and structure of fungal communities in soils containing elevated concentrations of radionuclides are still insufficient to identify clear patterns of changes in the composition and structure of communities caused by this type of pollution. Nevertheless, our findings of decreased abundance of *Mortierella* in radioactively contaminated soil compared to control (Figure 4C) are consistent with data obtained for soils of uranium mining sites **[5]**, however, another study noted that *Mortierella sp*. not only effectively reduces the bioavailability of heavy metals, but also significantly shortens the reclamation period **[75]**. The abundances of *Suillus* and *Tylospora* genera in soils contaminated with Pb were decreased **[64]**, similar to reduction observed under elevated activity concentrations of ^238^U and ^226^Ra at the contaminated site studied here. Despite the fact that, *Suillus luteus* has been to exhibit tolerance to polymetallic contamination of the soil **[76–78]**.

## Conclusion

The results of this study, conducted using ITS metabarcoding, revealed no differences in the structure of soil fungal communities between sites differing in the content of naturally occurring radionuclides. Additionally, no significant variations in diversity levels were observed due to contamination intensity or seasonal changes. Overall, fungal taxa resistant to heavy metals, abiotic and biotic stress and capable of forming symbiotic relationships with plants were more prevalent in the radioactively contaminated soils. Statistically significant differences were identified in the abundance of ectomycorrhizal and white rot fungi between the contaminated and reference soils. The community composition which has developed over more than half a century supports stable functioning in soils with elevated concentrations of natural radionuclides such as ^238^U and ^226^Ra.

## Supplementary Materials

Table S1: The significance levels of the differences for 32 ASV from the contaminated and reference sites using DESeq2 analysis.

## Author Contributions

Conceptualization, Anna Rybak; Data curation, Anna Rybak; Formal analysis, Anna Rybak; Funding acquisition, Anna Rybak; Investigation, Anna Rybak, Tatiana Maystrenko, Elena Rasova, Elena Belykh and Ilya Velegzhaninov; Methodology, Anna Rybak, Tatiana Maystrenko, Elena Belykh and Ilya Velegzhaninov; Project administration, Anna Rybak; Supervision, Anna Rybak; Visualization, Anna Rybak; Writing – original draft, Anna Rybak, Tatiana Maystrenko, Elena Rasova and Elena Belykh; Writing – review & editing, Anna Rybak, Tatiana Maystrenko, Elena Belykh and Ilya Velegzhaninov. All authors have read and agreed to the published version of the manuscript.

## Funding

This study was funded by a Russian Science Foundation Grant No. 23-74-01125, (https://rscf.ru/project/23-74-01125/).

## Institutional Review Board Statement

No applicable.

## Informed Consent Statement

No applicable.

## Data Availability Statement

The original contributions presented in this study are included in the article/supplementary material. Further inquiries can be directed to the corresponding author(s).

## Acknowledgments

The studies of biological activity were carried out using the equipment of the Center for Collective Use «Molecular Biology» at the Institute of Biology of the Komi Scientific Centre of the Ural Branch RAS.

## Conflicts of Interest

The authors declare no conflicts of interest.

## Notes

### Competing Interest Statement

The authors have declared no competing interest.

