## Supplemental Table S1 for "Fungal communities in long-term contaminated soils within a former radium production area": Supplementary Table S1.docx

**Table S1**. The significance levels of the differences for 32 ASV from the contaminated and reference sites using DESeq2 analysis

| **ASV** | **baseMean** | **logFC** | **lfcSE** | **stat** | **p-value** | **padj** | **Genus (Phylum)** |
| --- | --- | --- | --- | --- | --- | --- | --- |
| ASV15 | 535.606 | 28.719 | 2.635 | 10.899 | 1.17E-27 | 1.14E-24 | *Cortinarius* (Basidiomycota) |
| ASV146 | 51.786 | 25.320 | 2.776 | 9.122 | 7.40E-20 | 3.63E-17 | *Archaeorhizomyces* (Ascomycota) |
| ASV43 | 54.318 | -24.142 | 2.725 | -8.859 | 8.06E-19 | 2.64E-16 | *Russula* (Basidiomycota) |
| ASV131 | 70.034 | 25.931 | 2.957 | 8.769 | 1.80E-18 | 4.41E-16 | *Serendipita* (Basidiomycota) |
| ASV66 | 65.152 | 25.828 | 2.957 | 8.734 | 2.45E-18 | 4.80E-16 | *Amphinema* (Basidiomycota) |
| ASV88 | 57.282 | 25.643 | 2.957 | 8.672 | 4.26E-18 | 6.96E-16 | *Cortinarius* (Basidiomycota) |
| ASV83 | 51.071 | 25.500 | 2.957 | 8.623 | 6.51E-18 | 9.12E-16 | *Cortinarius* (Basidiomycota) |
| ASV13 | 39.012 | 25.135 | 2.957 | 8.499 | 1.91E-17 | 2.34E-15 | *Russula* (Basidiomycota) |
| ASV21 | 37.832 | 25.092 | 2.957 | 8.485 | 2.16E-17 | 2.36E-15 | *Cortinarius* (Basidiomycota) |
| ASV114 | 32.882 | 24.893 | 2.957 | 8.417 | 3.87E-17 | 3.79E-15 | *Clavaria* (Basidiomycota) |
| ASV117 | 31.802 | 24.844 | 2.957 | 8.400 | 4.46E-17 | 3.97E-15 | *Cortinarius* (Basidiomycota) |
| ASV62 | 29.264 | 24.739 | 2.958 | 8.365 | 6.02E-17 | 4.22E-15 | *Unclassified* (Ascomycota) |
| ASV63 | 85.705 | -24.777 | 2.957 | -8.379 | 5.32E-17 | 4.22E-15 | *Suillus* (Basidiomycota) |
| ASV124 | 30.555 | 24.751 | 2.958 | 8.369 | 5.81E-17 | 4.22E-15 | *Cortinarius* (Basidiomycota) |
| ASV236 | 25.030 | 24.518 | 2.958 | 8.289 | 1.14E-16 | 7.45E-15 | *Unclassified* (Fungi) |
| ASV50 | 22.983 | 24.412 | 2.958 | 8.253 | 1.54E-16 | 9.45E-15 | *Unclassified* (Basidiomycota) |
| ASV37 | 59.059 | -24.258 | 2.957 | -8.203 | 2.34E-16 | 1.35E-14 | *Tylospora* (Basidiomycota) |
| ASV51 | 55.065 | -24.156 | 2.957 | -8.169 | 3.11E-16 | 1.70E-14 | *Cortinarius* (Basidiomycota) |
| ASV95 | 41.316 | -23.759 | 2.957 | -8.035 | 9.40E-16 | 4.85E-14 | *Unclassified* (Basidiomycota) |
| ASV48 | 36.107 | -23.566 | 2.957 | -7.969 | 1.60E-15 | 7.84E-14 | *Unclassified* (Ascomycota) |
| ASV17 | 10.625 | 23.293 | 2.959 | 7.871 | 3.51E-15 | 1.64E-13 | *Cortinarius* (Basidiomycota) |
| ASV19 | 28.505 | -23.242 | 2.957 | -7.859 | 3.87E-15 | 1.68E-13 | *Hymenogaster* (Basidiomycota) |
| ASV99 | 28.455 | -23.237 | 2.957 | -7.857 | 3.93E-15 | 1.68E-13 | *Thelephora* (Basidiomycota) |
| ASV69 | 17.841 | -22.586 | 2.958 | -7.636 | 2.24E-14 | 9.17E-13 | *Mortierella* (Mortierellomycota) |
| ASV98 | 20.396 | -22.529 | 2.958 | -7.617 | 2.60E-14 | 1.02E-12 | *Hyaloscypha* (Ascomycota) |
| ASV109 | 5.149 | 22.376 | 2.962 | 7.554 | 4.21E-14 | 1.59E-12 | *Suillus* (Basidiomycota) |
| ASV65 | 30.951 | -22.299 | 2.957 | -7.540 | 4.69E-14 | 1.70E-12 | *Unclassified* (Ascomycota) |
| ASV55 | 11.987 | -22.034 | 2.958 | -7.448 | 9.49E-14 | 3.32E-12 | *Piloderma* (Basidiomycota) |
| ASV207 | 14.782 | 21.888 | 2.959 | 7.398 | 1.38E-13 | 4.66E-12 | *Russula* (Basidiomycota) |
| ASV158 | 10.597 | -21.864 | 2.959 | -7.390 | 1.47E-13 | 4.81E-12 | *Unclassified* (Ascomycota) |
| ASV140 | 10.793 | -21.799 | 2.959 | -7.368 | 1.73E-13 | 5.48E-12 | *Unclassified* (Basidiomycota) |
| ASV1263 | 1.412 | -18.947 | 2.973 | -6.373 | 1.85E-10 | 5.67E-09 | *Unclassified* (Fungi) |

Note: *baseMean* - mean of normalized counts for all samples. *logFC* - log2 fold change. *lfcSE* - standard error. *stat* - Wald statistic. *p-value* - Wald test p-value. *padj* - Benjamini and Hochberg method adjusted p-values.
